# Morphological and biochemical repercussions of *Toxoplasma gondii* infection in a 3D human brain neurospheres model

**DOI:** 10.1101/2020.08.31.274985

**Authors:** Paulo Emílio Corrêa Leite, Juliana de Araujo Portes, Mariana Rodrigues Pereira, Fabiele Baldino Russo, Erica S. Martins-Duarte, Nathalia Almeida dos Santos, Marcia Attias, Francisco J. Barrantes, Patricia Cristina Baleeiro Beltrão-Braga, Wanderley de Souza

**Author notes:** Corresponding authors, Av. Nossa Senhora das Graças 50, LABET - Dimav, Prédio 27, Duque de Caxias, Xerem, RJ, 25250-020, Brazil., Laboratório de Ultraestrutura Celular Hertha Meyer, Instituto de Biofísica Carlos Chagas Filho, CCS, UFRJ, Av. Carlos Chagas 373, CCS, Cidade Universitária, Rio de Janeiro, RJ, 21941-902, Brazil.

## Abstract

**Background:** Toxoplasmosis is caused by the parasite *Toxoplasma gondii* that can infect the central nervous system (CNS), promoting neuroinflammation, neuronal loss, neurotransmitter imbalance and behavioral alterations. *T*. *gondii* infection is also related to neuropsychiatric disorders such as schizophrenia. The pathogenicity and inflammatory response in rodents are different to the case of humans, compromising the correlation between the behavioral alterations and physiological modifications observed in the disease. In the present work we used BrainSpheres, a 3D CNS model derived from human pluripotent stem cells (iPSC), to investigate the morphological and biochemical repercussions of *T*. *gondii* infection in human neural cells.

**Methods:** We evaluated *T. gondii* ME49 strain proliferation and cyst formation in both 2D cultured human neural cells and BrainSpheres. Aspects of cell morphology, ultrastructure, viability, gene expression of neural phenotype markers, as well as secretion of inflammatory mediators were evaluated for 2 and 4 weeks post infection in BrainSpheres.

**Results:** *T. gondii* can infect BrainSpheres, proliferating and inducing cysts formation, neural cell death, alteration in neural gene expression and triggering the release of several inflammatory mediator.

**Conclusions:** BrainSpheres reproduce many aspects of *T. gondii* infection in human CNS, constituting a useful model to study the neurotoxicity and neuroinflammation mediated by the parasite. In addition, BrainSpheres can be an important tool for better understanding the possible correlation between psychiatric disorders and human CNS infection with *T. gondii*

## 1. INTRODUCTION

Toxoplasmosis is caused by an obligatory intracellular parasite, the protozoan *Toxoplasma gondii*, one of the world’s most common zoonoses that affects around one third of the human population (1). The majority of exposed individuals are asymptomatic and do not develop clinical manifestations of infection (2). The acute form of the disease evolves into a latent, chronic condition, with *T. gondii* localized inside tissue cysts persisting in infected individuals for life. *T. gondii* can cross the blood-brain barrier and infects the central nervous system (CNS), mainly the amygdala, hippocampus, olfactory bulb and cortex (3-5). The infection can disturb neural circuits, inducing neuroinflammation, neuronal loss and deficits in CNS functioning, indicating a possible correlation between *T. gondii* infection and neurodegenerative disorders, such as Alzheimer’s and Huntington diseases (6, 7). Moreover, prenatal exposure to *T. gondii* increases the probability of congenital syndromes, including CNS disturbances like severe intellectual disability and seizures in the phenotype of affected newborns (8).

The latent infection is also related to psychomotor deficits, a lower intelligence quotient and personality changes (9, 10), generating strong interest in elucidating how *T. gondii* infection affects human behavior (11). Toxoplasmosis has been associated with suicidal tendencies, traffic accidents, cognitive deficits, psychiatric disorders, anxiety, aggressiveness and violence (11-19). For individuals who attempted suicide, higher IgG antibody levels against *T. gondii* were reported, suggesting that parasite infection could enhance human behavioral alterations (20). Behavioral alterations have also been observed in infected rodents, who developed changes in predation risk as observed through their reduced fear of cats (21, 22), being later also reported for other predators (23). This effect could be related to the known tropism of *T. gondii* for the amygdala, a brain region involved in fear processing. This area shows increased cyst density and dendritic retraction in its basolateral portion after infection (24, 25). Additionally, infected rodents display impaired motor performance, learning and memory deficits (26, 27).

*T. gondii* infection has also been related to neuropsychiatric disorders such as schizophrenia spectrum disorders, bipolar disorder and anxiety (28-30); these reports are reinforced by the higher IgG levels to *T. gondii* found in schizophrenic patients than in control individuals (31, 32). The neuropsychiatric disorders observed in individuals infected by *T. gondii* may be related to neurotransmitter imbalance promoted by the parasite invasion in the CNS. The levels of dopamine, adrenaline, noradrenaline, serotonin, glutamate, GABA, acetylcholine and other neurotransmitters are affected by the infection (33-37), and their deregulation is already known to be related to mental disorders (38, 39). Also, rodents show an increase in dopamine release when infected by *T. gondii* (35). Dopamine is also secreted by the cysts in infected rodents and humans, contributing to the increase in local dopamine levels that may be associated with behavioral changes, as observed in schizophrenic patients (35, 40, 41). However, striatal dopamine imbalance may not be the only neurotransmitter alteration in schizophrenia spectrum disorders in infected individuals. Recent studies have shown the importance of neuroinflammation mediators in the behavioral alterations observed in *T. gondii* infection, a process also observed in neuropsychiatric disorders (23, 42). Behavioral alterations were observed in mice infected with a *T. gondii* strain incapable of inducing cyst formation, suggesting that neuroinflammation could be responsible (22, 42).

Neurotransmitter and neuroinflammation imbalance may thus play an important role in the behavioral alterations brought about by *T. gondii* infection in animal models and in humans. However, the precise nature of the correlation remains to be investigated. The characteristics of the disease in terms of pathogenicity and inflammatory response differ in humans compared to rodents. Animal models have provided a useful tool to study *T. gondii* infection, in particular to address the initial phase of host-parasite interaction at the cellular and biochemical levels.

Despite the usefulness of the data derived from experimental animal systems, these models fall short in addressing certain specific aspects of human cell physiology and kinetics. In this work, we used a 3D CNS model derived from human pluripotent stem cells (iPSC), BrainSpheres, to investigate morphological and biochemical repercussions of *T. gondii* infection with a view to elucidating the possible correlation of *T. gondii* infection with human behavioral alterations and neuropsychiatric disorders.

## 2. METHODS AND MATERIALS

### 2.1 iPSC generation

iPSCs were generated from Stem Cells from Human Exfoliated Deciduous Teeth (SHED), from control donors recruited through the Tooth Fairy Project initiative (University of São Paulo), with the approval of the Ethics Committee of the Institute of Biosciences, University of São Paulo (Protocol 1001). Cellular reprogramming experiments were conducted using the Sendai virus (CytoTune, Gibco, Life Technologies) carrying the four factors Oct4, Sox2, Kfl4 and c-Myc (43). Briefly, SHED were transduced, transferred two days later to a feeder layer condition, consisting of murine embryonic fibroblasts (MEFs, Millipore) and maintained in DMEM/F12 (Gibco, Life Technologies), 20% Knockout Serum Replacement (Invitrogen), 1% non-essential amino acids, and 100 μM beta-mercaptoethanol. The iPSCs colonies were identified after approximately 3 weeks in this culture system and subsequently transferred to Matrigel (BD Biosciences) coated plates. Once on feeder-free plates, iPSCs were maintained in mTeSR medium (Stem Cell Technologies), changed daily.

### 2.2 NPC generation

Following the fifth day of seeding iPSCs on matrigel, the medium was changed for 48 h to N2: DMEM/F12 medium supplemented with 1x N2 supplement (Invitrogen) and dual SMAD inhibitors 1 μM of dorsomorphin (Tocris) and 10 μM of SB431542 (Stemgent). Subsequently, colonies were manually scraped and cultured in suspension as Embryoid Bodies (EBs) for five days at 90 rpm in N2 medium. Next, EBs were plated on matrigel-coated plates with NGF media composed of: DMEM/F12 medium supplemented with 0.5x N2, 0.5x Gem21 supplement (Gemini Bioproducts), 20 ηg/mL FGF2 and 1% penicillin/ streptomycin. Emerged rosettes containing the neural progenitor cells (NPCs) were manually selected, dissociated and plated in a double-coated plate with poly-ornithine (10 μg/mL-Sigma) and Laminin (2.5 μg/mL, Thermo Fisher Scientific). This NPC obtained population was expanded using NGF medium.

### 2.3 Maintenance of ME49 Toxoplasma gondii strain

The *T. gondii* strain ME49-Luc, a type II strain engineered to express luciferase and GFP (44) (kindly provided by Dr. John C. Boothroyd, Stanford University), was used in this work. Me49-Luc tachyzoites were maintained *in vitro* in monolayers of Normal Human Dermal Fibroblasts-Neonatal (NHDF, Lonza; kindly provided by Dr. Sheila Nardelli - ICC/FIOCRUZ-BR) cultured in 25 cm^2^ flasks with RPMI 1640 medium (Gibco) supplemented with 2% FBS (Gibco), Pen/Strep/Fungizone (Gibco) and 2 mM glutamine (Sigma-Aldrich). Cultures were kept in an incubator at 37°C in a humid atmosphere with 5% CO_2_. The parasites were released from infected cells with a rubber scraper, followed by serial passages through needles of different gauges (22, 25 and 27G). Parasites were then separated from host cell debris by filtration through a 5 μm membrane. After filtration, parasites were centrifuged (2000 xg for 10 min) and the obtained pellet was then recovered in 1mL RPMI medium without antibiotics and serum. The number of parasites in 1 mL was quantified using a Neubauer chamber.

### 2.4 NPC maintenance, 2D neuronal culture and infection

NPC were differentiated from iPSC and maintained up to 12 passages in proliferation medium: KO DMEM/F12 medium supplemented with 1x StemPro supplement, 20 ng/mL human bFGF, 20 ng/mL epidermal growth factor (EGF), 4 mM L-glutamine, 500 Units Penicillin and 500 μg Streptomycin. All these products were purchased from ThermoFisher (USA). Half of the medium was replaced every 48 h.

NPC were mechanically detached at 90% confluence and resuspended in the differentiation medium: Neurobasal Electro medium supplemented with B-27-electro, 10 ng/mL BDNF and 10 ng/mL GDNF, 4 mM L-glutamine, 500 Units Penicillin, and 500 μg Streptomycin. All these products were purchased from ThermoFisher (USA). The cells were seeded on glass coverslips previously coated with 10 µg/mL poly-L-ornithine and 2.5 µg/mL laminin (both from Sigma Aldrich) for 12 h at 5×10^4^ cells per well in 24-well plates, to be later submitted to IF staining. Cells were maintained in a humidified incubator at 37°C and 5% CO_2_. The medium was replaced every 72 h. Cells were used for experiments after 4 weeks of differentiation. The *T. gondii* ME49-Luc strain was diluted in differentiation medium just before the infection at 1:100 parasite/cell proportion. The cells were infected after 4 weeks of differentiation (day 28). Cell samples were processed for endpoint measurements at the indicated time points. Bright field images were acquired in a Leica DMI3000 microscope (Wetzlar, Germany).

### 2.5 3D organotypic culture and infection

For BrainSpheres differentiation (see description in (45)), cells were mechanically detached upon reaching 90% confluence. The cells were seeded at 4×10^3^ cells per hydrogel hollow mold with 400 µm diameter made in the laboratory using the differentiation medium described above. Cells were placed into a humidified incubator at 37°C and 5% CO_2_. The medium was replaced every 72h. The BrainSpheres were used for experiments after 4 weeks of differentiation.

The ME49-Luc strain was diluted in the differentiation medium before the infection at 1:25 parasite/cell proportion. The BrainSpheres were infected after 4 weeks of differentiation (day 28). The supernatant or spheroid samples were collected for endpoint measurements at the indicated time points. The spheroid size was quantified in bright field micrograph images acquired in a Leica DMI3000 microscope using ImageJ software (https://imagej.nih.gov/ij/index.html, NIH).

### 2.6 Immunofluorescence assay

Samples were fixed and blocked with 2% BSA (Sigma-Aldrich) for 4 h, followed by overnight incubation at 4°C with the primary antibodies for MAP2 (ab5392, Abcam) at 1:1000 and SOX2 (Cell Signaling Technology) at 1:200. Next, cells were washed three times with DPBS and submitted to second round blocking (2% BSA for 1 h) at room temperature. Alexa Fluor 488 and 555 secondary antibodies (Life Technologies) were used at 1:500 for 1 h at room temperature, followed by sample mounting using Pro-long Gold Antifade Reagent with DAPI (Invitrogen). Images were obtained using 40X objectives on a Zeiss microscope (Carl Zeiss Microscopy GmbH, Jena, Germany).

### 2.7 Histological staining and confocal microscopy

BrainSpheres were fixed with 4% formaldehyde, freshly prepared from paraformaldehyde (PFA) for 1 h in PBS, washed 3 times with PBS and incubated for 2 h with blocking and permeabilization buffer (1% BSA, 5% goat serum, 0.15% saponin (Sigma Aldrich)). Samples were incubated overnight with *Dolichos biflorus* lectin conjugated to Tetramethylrhodamine-5-(and 6)-isothiocyanate (DBA-TRITC, Vector Lab) at 10 μg/mL diluted in blocking buffer at 4°C, followed by three washing steps. Then, samples were mounted on glass slides with Prolong Gold-antifade with Dapi reagent (Molecular Probes) for confocal microscopy. Z-stacks were acquired starting from at the top of the sample with ∼100-200 nm optical sections. Images were obtained using a Structured Illumination Microscopy (SIM) mode in a Zeiss ELYRA PS.1 (Zeiss) microscope. All images were acquired with identical time exposure and image settings.

### 2.8 Scanning electron microscopy (SEM)

Differentiated NPCs and BrainSpheres after 4 weeks of differentiation were fixed with 2.5% glutaraldehyde in 0.1 M sodium cacodylate buffer (pH 7.4) for 2 h at room temperature. After three washing steps with PBS, the samples were post-fixed with 1% osmium tetroxide in 0.1 M sodium cacodylate buffer for 90 min in the dark, followed by three washing steps with 0.1 M cacodylate buffer and then three washings in distilled water. The samples were gradually dehydrated in ethanol series and dried by the CO_2_ critical-point method in a CPD300 (Leica). Samples were mounted on aluminum stubs and coated in Balzers Apparatus. Micrographs were obtained with 25 kV in a FEI QUANTA 250 SEM (Thermo Fisher).

### 2.9 Transmission electron microscopy (TEM)

BrainSpheres were fixed with 2.5% glutaraldehyde in 0.1 M sodium cacodylate buffer (pH 7.4) for 2 h at RT. After three washing steps with sodium cacodylate buffer (pH 7.4), the samples were post-fixed with 1% osmium tetroxide, 5 mM calcium chloride and 1.25% potassium ferrocyanide in 0.1 M sodium cacodylate buffer for 1 h in the dark, followed by three washing steps with 0.1 M cacodylate buffer and distilled water. The BrainSpheres were gradually dehydrated with acetone and embedded in Epoxy resin (PolyBed812, Polysciences). Ultrathin sections from the surface or core of the BrainSpheres were obtained with a Leica C7 ultramicrotome, collected with copper grids and stained with uranyl acetate and lead citrate. Grids were then observed in a 120 kV in a FEI Tecnai Spirit TEM.

### 2.10 RNA extraction and quantitative real-time polymerase chain reaction (qRT-PCR)

Total RNA was extracted from 2D NPC cell culture after infection for 3 days, and from BrainSpheres after infection for 2 and 4 weeks, using RNeasy mini kit (Qiagen) according to manufacturer’s instructions. The amount and purity of total RNA were evaluated with a UV spectrophotometer (NanoDrop 2000, Thermo Fisher), by A260/280 and 260/230 ratios, considering the cut-off values equal or greater than 2.0 and 1.8, respectively. Extracted RNA samples were treated for 10 min at 37°C with RNase-free DNase I. First-strand synthesis of cDNA was performed using First-Strand cDNA synthesis kit (GE Healthcare), according to manufacturer’s instructions. The DNA amplification was performed using GoTaq qPCR Master Mix (Promega) and carried out on a Real-Time PCR System (StepOne; Applied Biosystems). The following conditions were used: initial holding stage at 95°C for 2 min followed by 40 cycles: denaturation at 95°C for 15 sec, annealing and extension at 60°C for 1 min, applying a melt curve (slow ramp of 0.3°C/20 sec to 95°C). Amplification was performed using specific primers for GAPDH (NM_001289746.1), MAP2 (NM_002374.3), GFAP (NM_002055.4), OLIG1 (NM_138983.2), Synaptophysin (NM_003179.2), vGLUT1 (NM_020309.3), GAD65 (NM_000818.2), TH (NM_199292), and TOXO (XM_002367038.2). Data were analyzed using the StepOne software (Applied Biosystems). For both control and treated samples, the GAPDH Ct values were subtracted from the gene Ct values to obtain the ΔCt value. ΔΔCt values were obtained according to ΔCt (treated samples) - ΔCt (control samples). The relative quantity was calculated according to 2^-ΔΔCt^.

### 2.11 Lactate dehydrogenase (LDH) release assay

LDH release was determined by the colorimetric CytoTox 96 Cytotoxicity Assay kit (Promega). After BrainSpheres infection, 20 µL of the supernatants were transferred to 96-well plates followed by the addition of 20 µL substrate solution. After 30 min of incubation in the dark, 20 µL of stop solution were added to each sample. Color development was proportional to the number of cells with the disrupted plasma membrane. Absorbance was measured at 490 nm. LDH positive control and substrate were used as internal control according to manufacturer’s instructions.

### 2.12 Analysis of multiple secreted mediators

Determination of cytokines, chemokines and growth factors secreted by BrainSphere cultures after ME49 infection was carried through Luminex (Austin TX, USA) xMAP magnetic technology for the following analytes: IL-1β, IL-1ra, IL-2, IL-4, IL-5, IL-6, IL-7, IL-8, IL-9, IL-10, IL-12 (p70), IL-13, IL-15, IL-17, eotaxin, bFGF, GCSF, GM-CSF, IFN-γ, IP-10, MCP-1 (MCAF), MIP-1α, MIP-1β, PDGF-BB, RANTES, TNFα and VEGF. Analysis was performed as previously described (46). Briefly, after calibration and validation of Bio-Plex Magpix (Bio-Rad), reagent reconstitution and standard curve preparation, magnetic beads were added to each well of the assay plate. Each step was preceded by washing steps using an automated Bio-Plex Pro wash station (Bio-Rad). Then, samples, standard and controls were added, followed by detection antibodies and streptavidin-PE. Finally, magnetic beads were re-suspended and read. The values detected in culture medium without spheres (background) were subtracted from the samples, allowing them to access the protein levels secreted by cultures.

### 2.13 Statistical analysis

GraphPad Prism 7 (GraphPad Software Inc., La Jolla, CA, USA) was used to calculate mean and standard errors of the assays. One- or two-way ANOVA tests and unpaired t test were applied as appropriate to obtain the statistical significance of means. The concentration of each secreted inflammatory mediator was quantified with xPONENT software version 4.2 (Biorad Laboratories Inc., Hercules, CA, USA). Differences were considered statistically significant at the 0.05 level of confidence.

## 3. Results

### Cyst formation and proliferation of ME49 strain of T. gondii in monolayers of neural cell cultures

BrainSpheres developed from human NPC express mature neurons (MAP2) after 4 and 8 weeks of differentiation, while NPC (SOX2) exhibit a slight reduction over time (Supplemental Fig. 1). BrainSpheres were differentiated for 4 weeks before infection with the ME49 strain of *T. gondii* and infected cultures were observed until 4 days post-infection (dpi). After 1 dpi, it was possible to observe the expected neural projections in both control and infected cultures, without apparent differences. However, at 4 dpi, we observed structures with the characteristics of *T. gondii* cysts (Fig 1A, *arrowheads*).

**Fig 1.**
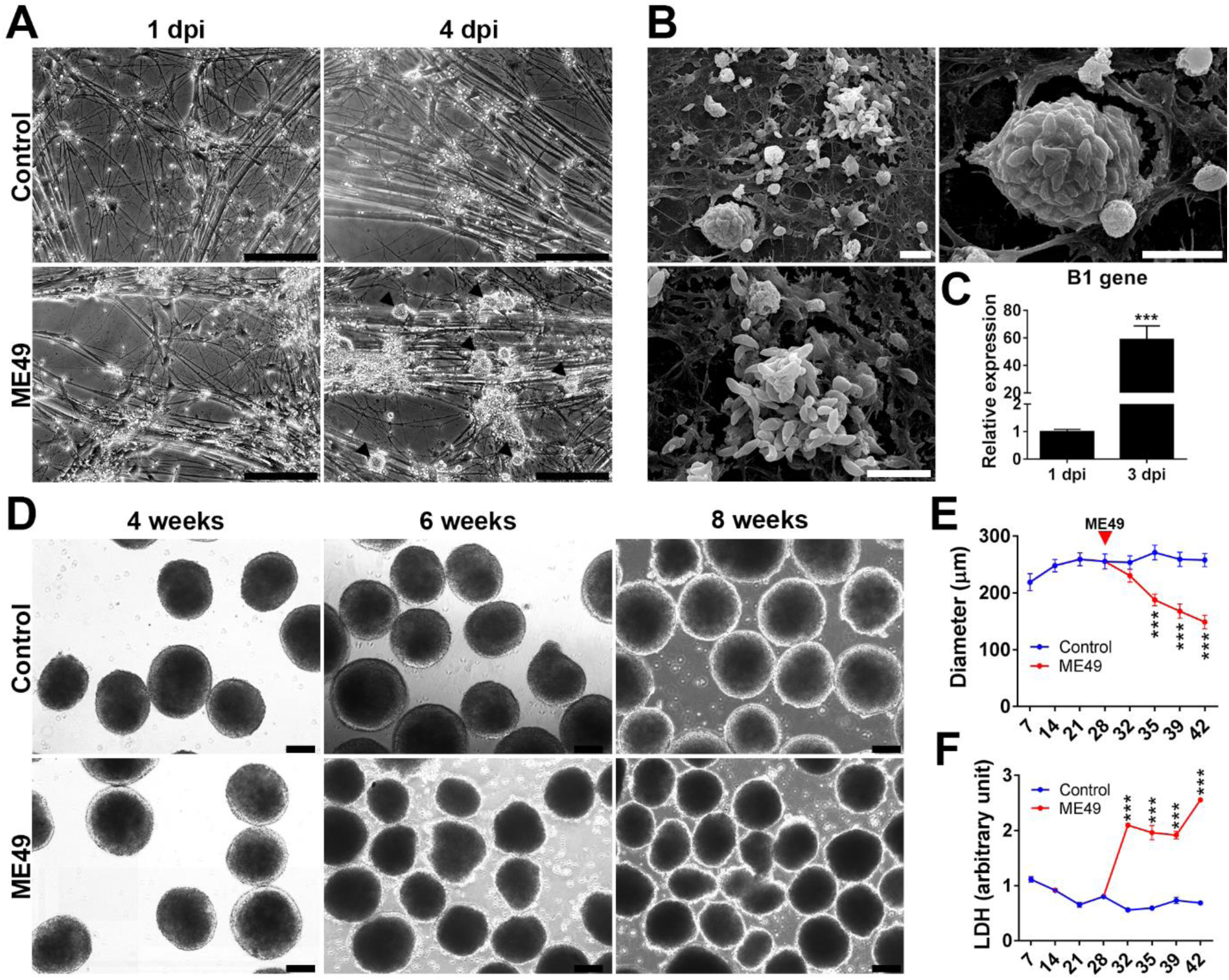
Cyst formation and proliferation of ME49 strain in monolayers of neural cell cultures and analysis of cell death in BrainSpheres. (A) Phase contrast images of monolayer neural cells differentiated during 4 weeks without infection (Control) and after 1 and 4 dpi with ME49 strain of *T. gondii*. Black arrowheads indicate cyst-like structures. (B) SEM micrographs showing an intact cyst and parasites from a disrupted cyst spreading to neighbor cells of monolayers of neural cell cultures. (C) qRT-PCR of *B1* gene of *T. gondii* after 1 dpi and 3 dpi of NPC cultures. (D) Bright field images of BrainSpheres differentiated for 4, 6 and 8 weeks without infection (Control) and after infection when BrainSpheres reach 4 weeks (day 28) of differentiation (ME49). (E) Quantification of BrainSpheres diameter during differentiation from day 7 to 42 (1 to 6 weeks of differentiation). Blue line corresponds to control non-infected and red line corresponds to ME49-infected. Red arrowhead indicates the moment of infection with ME49 strain of *T. gondii* at day 28. (F) Quantification of LDH in the BrainSpheres supernatant during differentiation from day 7 to 42. Blue line corresponds to control non-infected and red line to ME49-infected. Scale bars: (A) 100 µm; (B) 10 µm; (D) 100 µm. All data were collected from three independent experiments, with three technical replicates representing fold changes (± SD) for (C) and mean (± SEM) for (E) and (F). Student *t*-test was used to analyze the statistical significance for (C) and two-way ANOVA with Bonferroni’s post-test for (E) and (F) (\*\*\**p* < 0.001).

SEM showed the presence of intact and disrupted cysts at 4 dpi. It was also possible to observe the parasites emerging from the infected cells and spreading to the neighbor NPC in differentiation (Fig 1B). qRT-PCR analysis using primers for the *T. gondii* B1 gene showed that the *T. gondii* was able to proliferate, as the transcripts to B1 gene increased ∼60x after 3 dpi (*p* < 0.001) (Fig 1C).

### ME49 strain of T. gondii induces neural death in BrainSpheres

BrainSpheres are highly reproducible in size (200-260 μm) from batch to batch and experiment to experiment. Due to their small size, they do not develop a necrotic core as many larger sized organotypic models do (2–6), making this model very suitable for neural infection studies. BrainSpheres were infected when they reached 4 weeks (28 days) of differentiation. After this, infected and non-infected BrainSpheres were observed for a further 2 and 4 weeks (6 and 8 weeks of differentiation, respectively). Control BrainSpheres of all time points showed spherical shape and ∼250 µm in diameter from day 28 to 42 (6-weeks of differentiation) (Fig 1D, E). However, the infected BrainSpheres showed irregular shapes and a visibly reduced diameter from 6 to 8 weeks of differentiation (Fig 1D, E). Quantification of infected BrainSpheres showed average diameter of 150 ± 12 µm at day 42, corresponding to a 42% reduction compared to control samples (± 7%, *p* < 0.001) (Fig 1E).

LDH release, a marker of plasma membrane damage and cell death, was measured in infected and non-infected BrainSpheres. At day 32, after just 4 dpi of 4 weeks (day 28) of differentiation, ME49 strain of *T. gondii* infection of BrainSpheres sufficed to induce a large increase in LDH leakage (*p* < 0.001), reaching maximum levels at 42 dpi (6 weeks of differentiation) (*p* < 0.001) (Fig 1F).

### Morphological alterations and cystogenesis of ME49 strain of T. gondii in BrainSpheres

SEM shows the structural organization of the BrainSpheres after 4 weeks of differentiation. Uninfected BrainSpheres showed typical morphology with viable appearance (Fig 2A, C, E). However, infected BrainSpheres showed altered morphology and degenerated cells on the BrainSpheres surface (Fig 2B). Besides, it was possible to observe a considerable number of parasites inside the core of infected BrainSpheres (Fig 2D and F).

**Fig 2.**
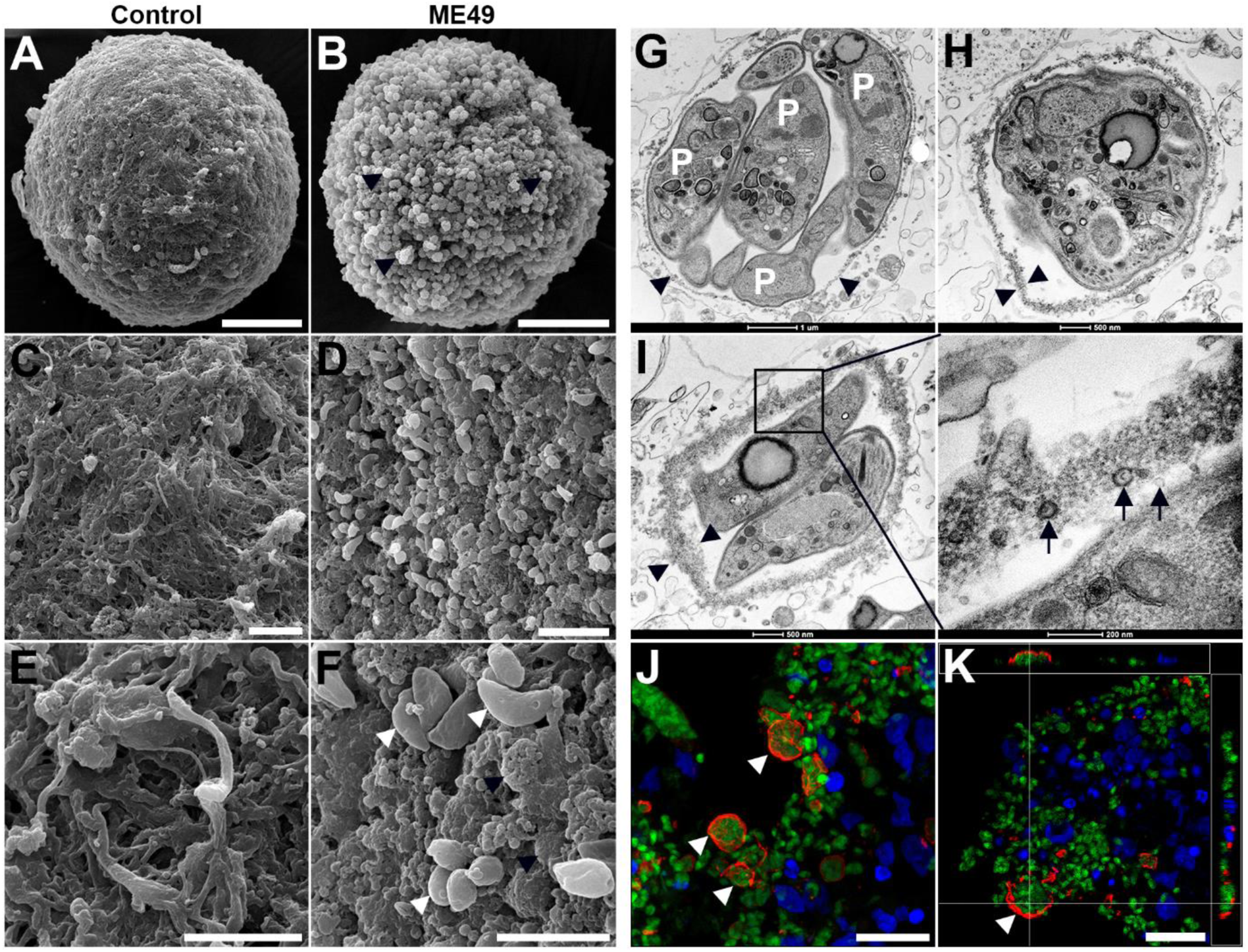
SEM, TEM and confocal microscopy images showing cellular damage and cystogenesis after infection of 4 weeks differentiated BrainSpheres with *T. gondii* from ME49-Luc (expressing luciferase and GFP) for 2 weeks (6 weeks total). (A, C, E) BrainSpheres maintained without infection (Control), showing (C, E) structures similar to neural projections. (B, D, F) SEM images of BrainSphere infected with *T. gondii*, it is possible to observe a pattern of generalized degeneration of the cells on the sphere surface (arrowheads in B). (D, F) A BrainSphere was fractured allowing the detection of several parasites (white arrowheads in F) and degenerated cells (black arrowheads in F) inside the 3D sphere. (G-I) It is possible to observe by MET observations the deposition of a dense material under the membrane of the parasitophorous vacuole, which leads to a gradual thickening (arrowheads). This indicates the formation of the cystic wall of bradyzoites. (inset in I) The cystic wall shows a particular composition with secreted materials by the parasites (arrows). P = parasites. (J, K) Confocal laser-scanning microscopy analysis after labeling infected BrainSpheres with *Dolichos biflorus* lectin (DBA-TRITC) (red) confirmed the presence of *T. gondii* cysts (arrowheads), and (K) parasites evading a disrupted cyst. The parasites from ME49-Luc show green fluorescence, DAPI labeled cell nuclei represented by blue fluorescence. Scale bars: (A, B) 50 µm; (C, D) 10 µm; (E, F) 5 µm; (G) 1 µm; (H, I) 500 nm; (inset in I) 200 nm; (J, K) 20 µm.

Analysis of BrainSpheres after 2 weeks of infection (6 weeks of differentiation) by TEM showed the deposition of a dense material under the membrane of the parasitophorous vacuole, leading to a gradual membrane thickening. The thickening of *T. gondii* parasitophorous membrane is indicative of cystic wall formation (Fig 2G-I) and, consequently, cystogenesis. The presence of cyst wall formation was confirmed by labelling the samples with DBA lectin, which specifically binds N-acetylgalactosamine residues at the bradyzoite cyst wall (47). The BrainSpheres showed a large number of parasitophorous vacuoles labeled with DBA, indicating cystogenesis due to differentiation of *T. gondii* tachyzoites to bradyzoite, the stage found during the chronic phase of toxoplasmosis (Fig 2J, K). Also, it is possible to observe parasites evading a disrupted cyst (Fig 2K).

### ME49 strain of T. gondii location at the periphery and in the core of BrainSpheres

Ultrastructural analysis of the BrainSphere subcellular morphology along the period of differentiation showed, as expected, the maintenance of their features and viability throughout the assays (Fig 3A, B). The 3D model shows the progression of the infection with the ME49 strain of *T. gondii* for at least 4 weeks. After 1 week of infection, it was possible to observe that the cells in the core of BrainSpheres contained several parasitophorous vacuoles with parasites showing numerous amylopectin-like granules in the cytoplasm, which is a characteristic of the bradyzoite stage. The development of bradyzoites was observed mostly in the core of the BrainSpheres (Fig. 3C-E).

**Fig 3.**
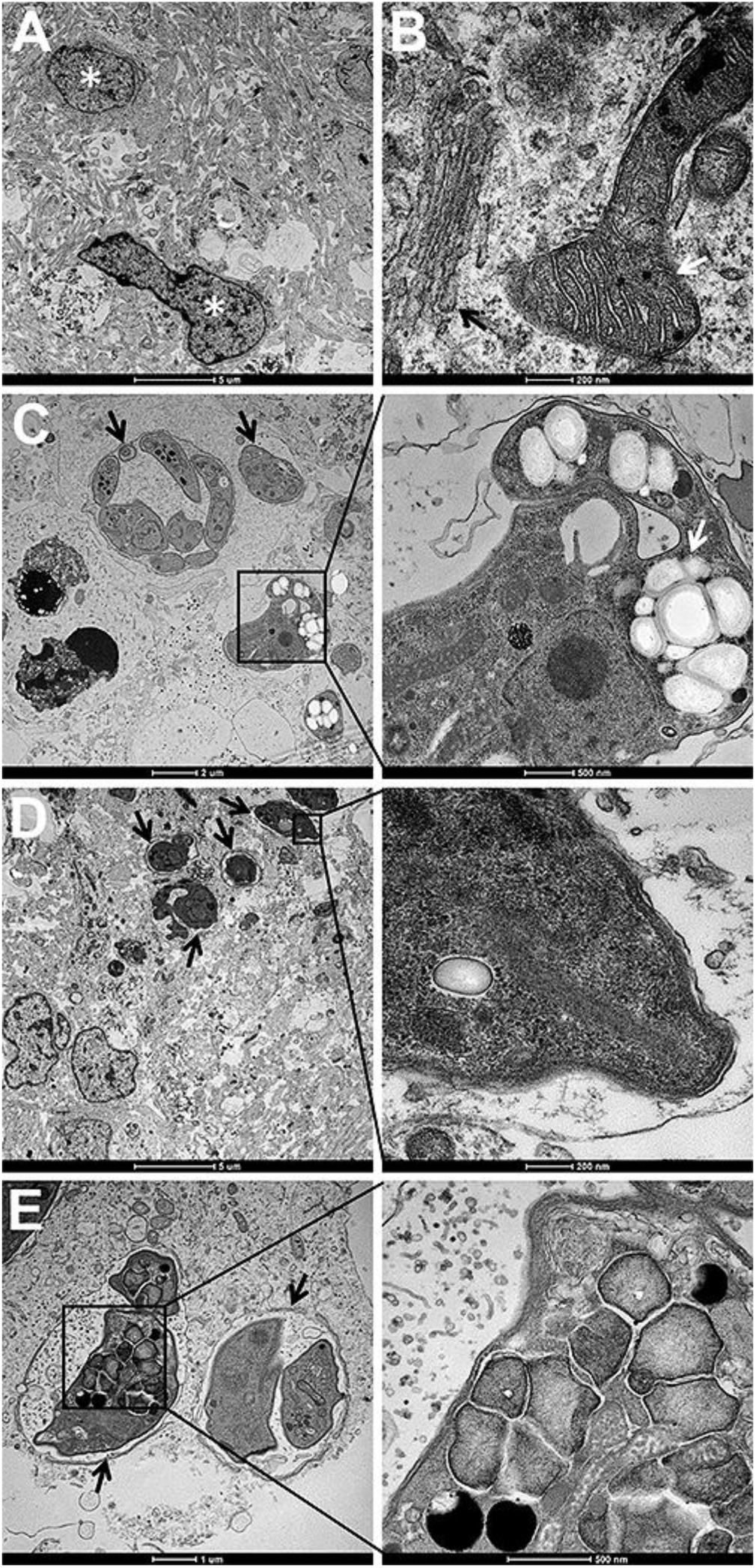
TEM images showing the core of BrainSpheres infected with ME49 strain of *T. gondii*. (A, B) BrainSpheres differentiated for 4 weeks and maintained without infection (Control) for another week. The BrainSpheres were accessed and show neural cells with typical cytoplasmic organization and cellular nuclei (asterisks in A). Indeed, organelles such as mitochondria (white arrow in B) and Golgi (black arrow in B) were observed. (C-E) After 4 weeks of differentiation, BrainSpheres were infected with *T. gondii* and then cultured for another week. Infected cells show parasitophorous vacuoles containing several parasites (black arrows) with preserved structures. The presence of parasites harboring granules similar to amylopectin confirms the differentiation process of tachyzoites to bradyzoites (white arrow inset in C). Scale bars: (A) 2 µm; (B) 200 nm; (C) 2 µm, inset 500 nm; (D) 5 µm, inset 200 nm; (E) 1 µm, inset 500 nm.

### ME49 strain of T. gondii alters the gene expression and myelination of neural cells in BrainSpheres

In order to explore how *T. gondii* infection affects the cell types, synapses and neuronal phenotypes in BrainSpheres, some specific mRNA levels were analyzed at different stages of differentiation and infection. The parasite tightly reduced the mRNA levels of MAP2 and GFAP, neuronal and astrocyte markers, respectively (Fig 4A). Indeed, levels of synaptophysin mRNA, a synapse marker, were also lower than in non-infected BrainSpheres, suggesting synapse loss. Interestingly, the synaptophysin mRNA levels were stable from the 4^th^ week (without infection) until the eighth week of differentiation (with infection) (Fig 4A). OLIG1 mRNA levels, an oligodendrocyte marker, were drastically increased soon after the *T. gondii* infection, mainly in the sixth week of differentiation, which corresponds to the second week of infection (Fig 4A). The infection induced a sharp increase in vGLUT1 and GAD65 mRNA levels (glutamatergic and GABAergic neuron markers, respectively), while tyrosine hydroxylase (TH) mRNA levels (dopaminergic neuron marker) sharply reduced, mainly in the sixth week of differentiation (Fig 4A). These results indicate that *T. gondii* infection differentially affects the neuronal phenotypes in BrainSpheres.

**Fig 4.**
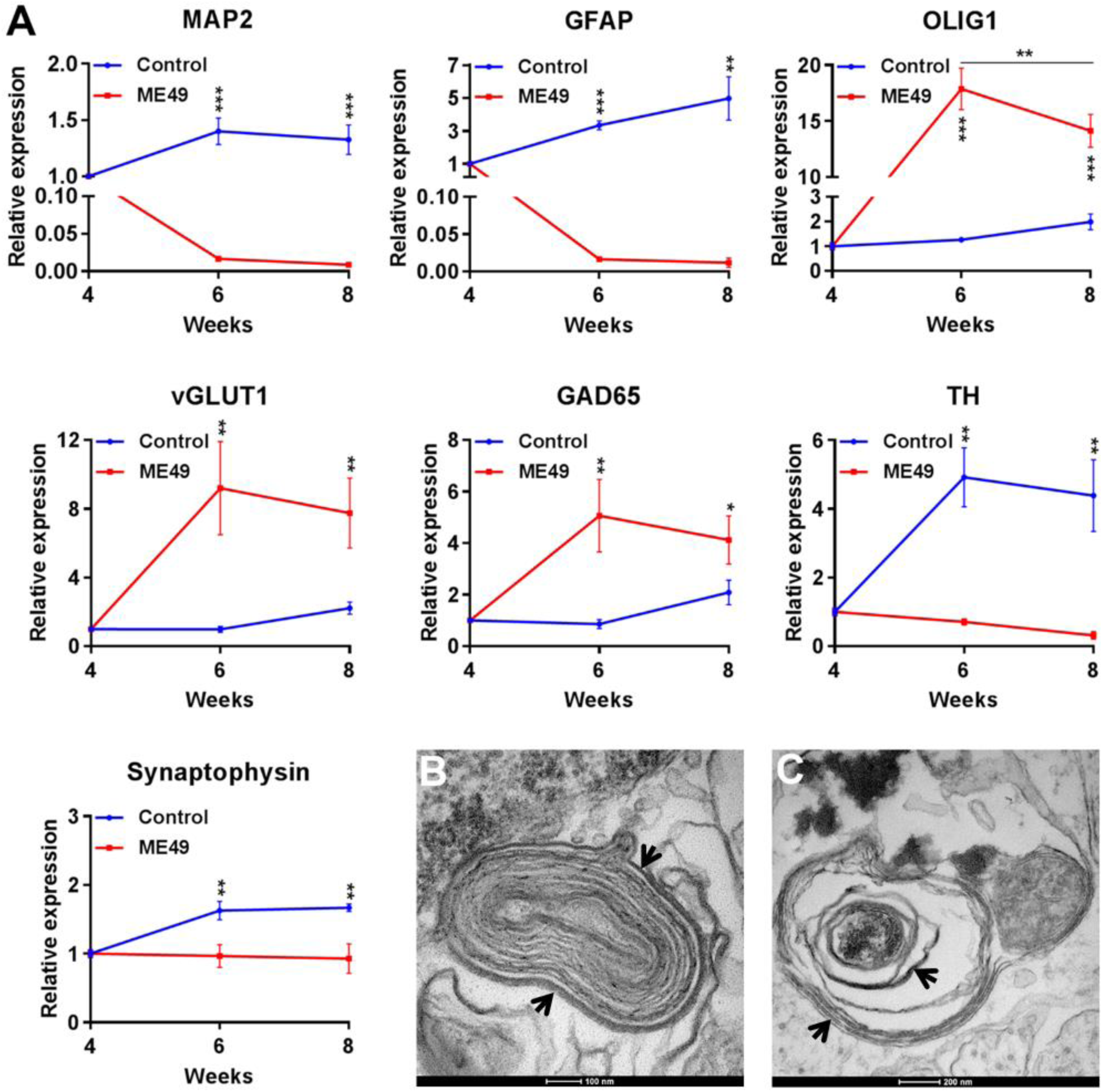
ME49 strain of *T. gondii* affects mRNA levels of cell types, synapses and neuronal phenotypes in BrainSpheres. (A) 3D human CNS model differentiated for 4, 6 and 8 weeks without infection (Control, blue line) and with infection (ME49, red line). The mRNA levels of different neural markers were analyzed: MAP2 (neuron), GFAP (astrocyte), OLIG1 (oligodendrocyte), synaptophysin (synapse), and vGLUT1, GAD65 and TH (glutamatergic, GABAergic and dopaminergic neurons, respectively). (B, C) TEM images of BrainSpheres differentiated for 4 weeks followed by *T. gondii* infection and cultured for 2 weeks. BrainSpheres without infection show a myelin-like pattern with normal concentric disposure (arrows, B), whereas infected BrainSpheres show a myelin loss pattern (arrows, C). Scale bars: (A) 100 nm; (B) 200 nm. Data were collected from three independent experiments and represent mean (± SD). Student *t*-test and two-way ANOVA with Bonferroni’s post-test were used to analyze the statistical significance (\**p* < 0.05, \*\**p* < 0.01, \*\*\**p* < 0.001).

Ultrastructural observations showed differences between the pattern of myelin-like structures in uninfected and infected BrainSpheres (Fig 4B, C). While in non-infected cells the structures show a normal morphology with concentric arrangement (Fig 4B), infection by *T. gondii* causes a remarkable alteration similar to myelin decompaction (Fig 4C).

### ME49 strain of T. gondii induces the release of cytokines, chemokines and growth factors in BrainSpheres

Analysis of multiple secreted proteins from non-infected BrainSpheres showed that most analytes were secreted in the initial phases of the differentiation (7, 14 and 21 days), with the levels decreasing over time. Despite the reduction in TNFα and IL-1β, these two cytokines showed a second small wave of increase from ∼28 to 42 days (Fig 5). Neuronal and glial cells produce lower levels of chemokines and cytokines than immune cells, but such production is critical to maintaining the homeostasis and microenvironment (48).

**Fig 5.**
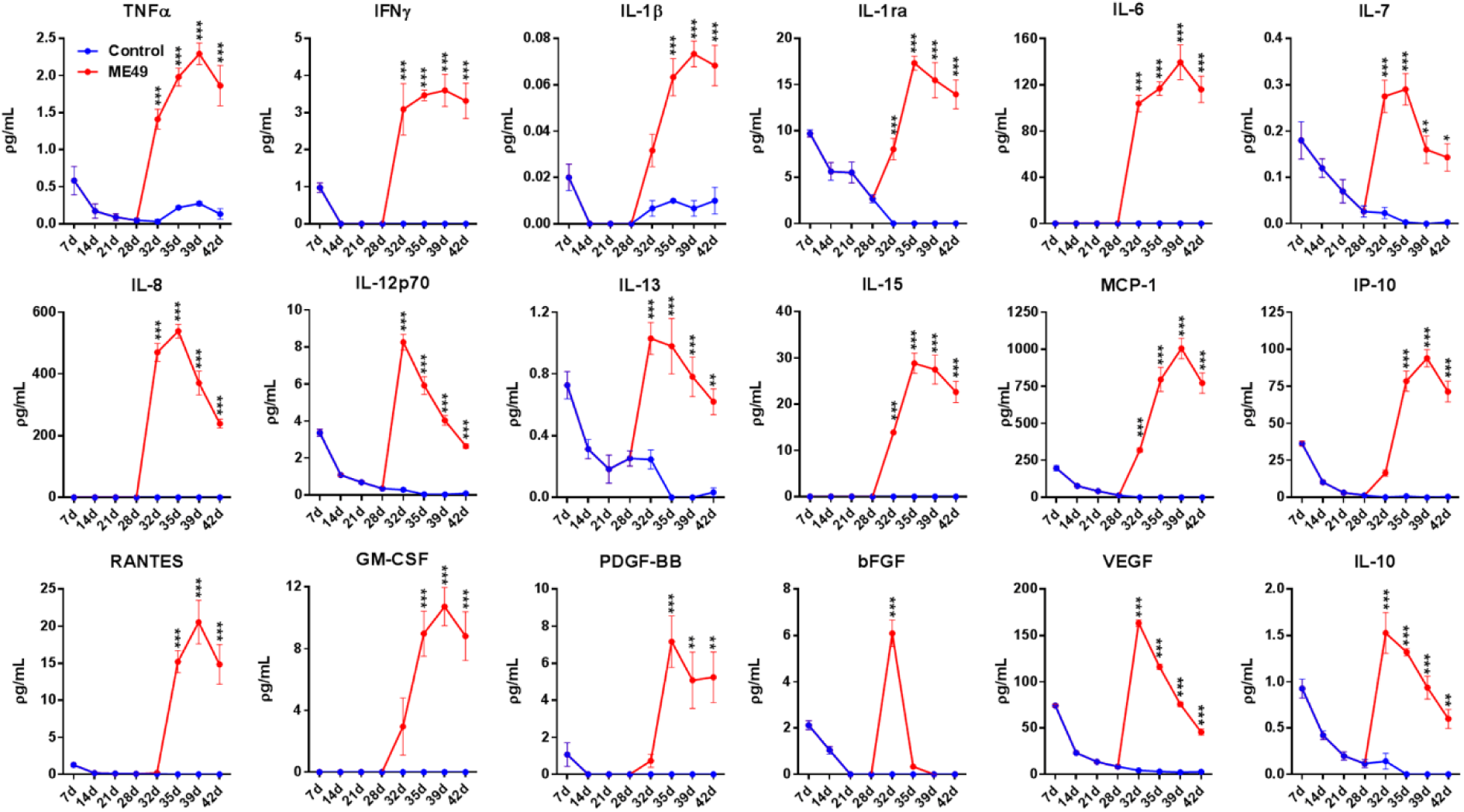
ME49 strain of *T. gondii* induces the release of cytokines, chemokines and growth factors in BrainSpheres. Graphs showing the levels of different secreted mediators of BrainSpheres without infection (Control, blue line) and after infection (ME49, red line) in different periods of BrainSpheres culture. The infection with ME49 strain of *T. gondii* occurred at day 28. The protein concentration levels of TNFα, IFNγ, IL-1β, IL-1ra, IL-6, IL-7, IL-8, IL-12p70, IL-13, IL-15, MCP-1, IP-10, RANTES, GM-CSF, PDGF-BB, bFGF, VEGF and IL-10 were analyzed. Data were collected from three independent experiments and represent mean (± SEM). Two-way ANOVA with Bonferroni’s post-test was used to analyze the statistical significance (\**p* < 0.05, \*\**p* < 0.01, \*\*\**p* < 0.001).

BrainSpheres infected with *T. gondii* evidenced a sharp increase in all shown analytes. The levels of potent pro-inflammatory cytokines like TNFα, IFNγ, IL-1β and IL-6 showed a sharp increase and a plateau up to 42 days of BrainSpheres culture. IL-15, which plays an important role in innate and adaptive immunity following infection, also seems to be maintained during the period of culture. Other pro-inflammatory cytokines like IL-7, IL-8, IL-12 and IL-13 showed a quick response soon after infection that persisted for 7 days (up to 35 days) and then started to drop. Chemokines displayed a pattern similar to that of the potent pro-inflammatory cytokines, but with a delayed response: IP-10, RANTES and GM-CSF showed no measurable levels until at least 4 days after infection at 32 days of culture. In contrast, MCP-1 showed significantly increased levels soon after infection (\*\*\**p* < 0.001). Growth factors such as GM-CSF and PDGF-BB also showed a delayed response, increasing their levels only after 7 days of infection at day 35 of culture. bFGF and VEGF showed a sharp increase immediately after infection at 32 days, but bFGF was almost undetectable at day 35 in culture, whereas VEGF showed a strong but gradual decrease. The anti-inflammatory mediators IL-1ra and IL-10 showed a sharp increase; however, while IL-1ra was maintained over time, IL-10 showed a strong but gradual decrease (Fig 5).

## 4. Discussion

Though rodent *in vivo* models are important tools for the analysis of various aspects of *T. gondii* infection, some of the results are not reproducible in humans due to differences between the species, highlighting the need for new models able to more closely represent human infection. Recently, our group established that BrainSpheres could be an useful tool to study the toxicity and biocompatibility of different nanomaterials for brain drug delivery (46). Compared to *in vitro* 2D cell culture, the 3D model displays more cell interactions and can better reproduce the *in vivo* physiology (49, 50). Thus, in this work, we used BrainSpheres as a 3D human CNS model to study the repercussions of *T. gondii* infection in the human brain.

*T. gondii* from avirulent strain ME49 was able to infect, proliferate and convert to cyst forms in both 2D human neural cell cultures and in the surface and deeper layers of the BrainSpheres. Both cysts and tachyzoites were found in the core of the BrainSpheres, indicating the invasiveness of *T. gondii* in the 3D model. Conversion to cyst forms was observed and the cystic wall of bradyzoites was confirmed by labelling with DBA lectin (47) and structural markers such as amylopectin granules and (51) deposition of a dense granular material under PV membrane. Cystogenesis is an important event related to the presence and permanence of *T. gondii* in the CNS, process mostly studied in rodents and 2D *in vitro* models (23, 52, 53).

In BrainSpheres, we observed that *T. gondii* infection causes a significant reduction in neural cells viability. A significant effect of infection was the reduction in size of the spheroids (∼42%), which correlated with the augmented neural cell death. Extensive damage to BrainSpheres was also evidenced by LDH leakage assay. Since the CNS is a site of high infection load, an important clinical manifestation of toxoplasmosis in reactivated patients is encephalitis (54, 55). In addition, focal brain necrosis is frequently observed in congenitally infected newborns. Thus, the cellular damage observed in infected BrainSpheres could result from the natural parasite replication cycle, causing cell lysis (56).

BrainSpheres expressed neuronal and glial markers, with distinct neuronal phenotypes similar to those observed in other studies (45, 50). We observed decreased mRNA levels of neuronal and astrocytic markers, consistent with the cell death findings. In contrast, we also observed increased OLIG1 mRNA levels, and alterations resembling myelin decompaction. We have not observed a predilection of *T gondii* infection for any particular cell type, concurring with *in vitro* studies using rodent and human cells that show cyst formation in both neurons and astrocytes (57, 58). It is known that cysts in neuronal processes usually result in neurite and synapse loss (52, 59). This is in agreement with our data showing that *T. gondii* infection reduced the synaptophysin mRNA levels compared to non-infected BrainSpheres, which can be interpreted as a reduction in synapse density. Neurite and synapse loss are observed in psychiatric disorders such as schizophrenia spectrum disorders and may be another manifestation of the correlation between *T. gondii* infection and mental disorders (60).

Changes in neuronal communication due to alterations in neurotransmitter release are an important consequence of the infection mediated by *T. gondii*. Our data discloses increased mRNA levels of vGLUT1 and GAD65 in the infected BrainSpheres, suggesting that the levels of the excitatory neurotransmitter glutamate and the inhibitory neurotransmitter GABA may increase as a consequence of *T. gondii* infection. It has previously been shown that extracellular glutamate levels were increased with *in vivo* infection, and the glutamate reuptake system mediated by astrocytes was affected as evidenced by a decrease in GLT-1 (glutamate transporter) levels (34). However, some proteins of glutamatergic synapse such as AMPA and NMDA receptor subunits were downregulated after *T. gondii* infection, suggesting reduced neurotransmission mediated by glutamate (61). Another *in vivo* study showed that *T. gondii* infection did not alter GAD67 protein levels but induced its mislocalization in neurons and the infected animals became more susceptible to seizures (33). It is possible that the increased GAD65 mRNA levels in infected BrainSpheres constitute a compensatory mechanism for its mislocalization, reflecting the attempt to restore the excitatory/inhibitory homeostasis.

Our data also evidenced alterations in another neurotransmitter system: the infection reduced TH mRNA levels in the BrainSpheres, suggesting the death of dopaminergic neuronal cells. Several works investigated *T. gondii* infection effects on dopamine levels, with many describing an increase in its levels. An *in vitro* study reported the reduction of TH protein levels in the presence of *T. gondii* without alteration of dopamine levels (52). Other authors did not observe changes in TH mRNA and protein levels, but instead reported increases in dopamine synthesis in PC12 cells (35). Similarly, augmented dopamine levels were found in infected rodents (62). Parasite cysts in rat cortical neuron cultures and mouse brains also display high dopamine levels. *T. gondii* expresses endogenous TH that synthetizes dopamine, thus additionally contributing to the augmented neurotransmitter levels (63). Dopamine neurotransmission is involved in motor control, cognition and emotion, and dysregulated levels of this neurotransmitter could be related to the behavioral alterations observed in *T. gondii* infection. In addition, dopamine dysfunction is associated with neuropsychiatric disorders like schizophrenia spectrum disorders (40, 64).

Nicotinic cholinergic neurotransmission may also play a role in schizophrenia spectrum disorders and other psychiatric illnesses (64). The enteric system’s infection produces the death of cholinergic neurons in the submucosal plexus of the duodenum-ileum (65), indicating a possible tropism of the parasite for the cholinergic system. Another possible involvement of the cholinergic system in CNS pathology in toxoplasma infection is the astrocyte-derived tryptophan metabolite, kynurenic acid. Electrophysiological experiments indicate that together with kynurenic acid, nicotinic acetylcholine receptors regulate the GABAergic inputs to CA1 pyramidal neurons in the hippocampus (66), with kynurenic acid acting as an antagonist of α7 nicotinic receptors (67). Although, this contention is still controversial. Instead, kynurenic acid is more likely to antagonize N-methyl-D-aspartate (NMDA) receptors (68). In addition, neuroinflammation increases kynurenic acid synthesis in CNS and this effect may contribute to the glutamatergic, GABAergic and cholinergic system modulations observed in *T. gondii* infection. Animal and human *in vivo* studies suggest that increased levels of kynurenic acid can be related to cognitive deficits observed in schizophrenia spectrum disorders (69). Increased production of kynurenic acid mediated by neuroinflammation may thus be another connection between *T. gondii* infection and neuropsychiatric disorders.

BrainSpheres exposed to *T. gondii* display increased release levels of several inflammatory mediators such as TNFα, IFNγ, IL-1β, IL-6, chemokines and growth factors. Astrocytes and microglia are activated and release inflammatory mediators during the CNS infection, including that mediated by *T. gondii* (5, 70*). As microglia is not present in our 3D human CNS model, these mediators are released by astrocytes and neurons; both cell types are known to release inflammatory mediators in response to T. gondii* (71). Infected mice cerebellar granule neurons in culture produce IL-6, TGF-β1, MIP-1α and MIP-1β, while astrocytes release IL-1α, IL-6, GM-CSF, MCP-1 and IP-10 (57, 70, 71). In addition, the CNS of infected animals displayed increased levels of TNF-α, IFNγ and IL-6 (72), which is following our data using the BrainSpheres model. IL-4 and IL-13 regulate the induction of indoleamine 2,3-dioxygenase activity that affects tryptophan availability, causing starvation and controlling *T. gondii* replication in human fibroblasts activated by IFNγ (73). A recent study showed that infected mice have sustained inflammation even during chronic infection, and several inflammatory mediators are still upregulated after the parasite clearance (23). In mice, some cytokines like IFNγ and TNFα are important to *T. gondii* infection control (74) and their signaling deficiency may be related to severe and reactivated cerebral toxoplasmosis in humans. However, only two patients of 180 cases of toxoplasmosis showed congenital immunodeficiency, corroborating the low reporting of patients with severe toxoplasmosis and primary immunodeficiency (75).

Anti-inflammatory cytokines are also crucial to control the *T. gondii* inflammation. For instance, IL-10 produced by monocytes contribute to the regulation of cerebral toxoplasmosis (76). Besides, IL-10 decreases animal mortality promoted by *T. gondii* and downregulates the immune response in chronic cerebral toxoplasmosis (77, 78). In BrainSpheres, IL-10 showed a sharp increase just after infection, but its levels gradually receded over time. Meanwhile, IL-1ra that binds to IL-1 receptor in cell membranes preventing IL-1 downstream signaling and activation of inflammation (79), was maintained over time. These data show that IL-10 and IL-1ra are responsive to *T. gondii* infection and may attempt to control the increased pro-inflammatory mediators.

Neuroinflammation mediated by *T. gondii* also promotes chemokines and growth factors release. Infected astrocytes in culture showed increased levels of the MCP-1 involved in CNS leukocyte infiltration (80); its receptor CCR2 also plays a relevant role in controlling CNS *T. gondii* infection (81). Mice with toxoplasmic encephalitis display astrocytes producing IP-10, critical for effector T cell trafficking and host survival in *T. gondii* infection, and MCP-1 (70, 82). Previously, it was shown that PDGF released from human platelets activated by *T. gondii* infection inhibited the intracellular parasite growth in human pulmonary fibroblasts (83), and that VEGF can control the *T. gondii* proliferation in human retinal pigment epithelium (84). Besides, GM-CSF levels were increased in cultured astrocytes and mice brain infected with *T. gondii* (85, 86). It is therefore possible that the increased levels of chemokines and growth factors in infected BrainSpheres are an attempt to maintain cell survival.

The behavioral alterations mediated by *T. gondii* infection are also impacted by neuroinflammation. Post-mortem human brain analyses showed that behavioral alterations correlate with high amounts of parasite DNA in the CNS (87). However, the correlation between behavioral alterations and cysts presence in specific regions of CNS is still controversial. While some studies suggest that there is a possible connection, other findings emphasize the importance of neuroinflammation more than cyst load or location for the altered behavior (23, 42, 53). The reduction of cyst load in infected mice reversed the hyperactivity, but the regular locomotor activity was restored only through neuroinflammation reduction. It therefore appears that some behavioral alterations are more related to neuroinflammation than to a parasite effect (42). In this sense, the authors of an *in vivo* study suggested that neuroinflammation is the major factor for behavioral alterations (23). Neuroinflammation is also present in several neuropsychiatric disorders such as schizophrenia, whose behavioral symptoms could be related to inflammation-mediated by *T. gondii* infection. Antipsychotic drugs with activity against *T. gondii* decreased the behavioral symptoms and inflammation in schizophrenic patients (88).

Our experimental findings are in agreement with the literature data reinforcing the possible connection between *T. gondii* infection of human CNS, alterations in neurotransmitter systems, and neuropsychiatric diseases, particularly schizophrenia spectrum disorders, with important behavioral dysfunction, a subject worth exploring in further depth. In addition, in the present study, we show that BrainSpheres constitute a relevant tool to characterize the infection mediated by *T. gondii* in human neural tissue in a species-specific manner, thus contributing to determining a possible correlation between infection by this parasite and neuropsychiatric disorders.

## ACKNOWLEDGMENTS

This study was supported by the Coordenação de Aperfeiçoamento de Pessoal de Nível Superior - Brasil (CAPES) - Finance Code 001, and FAPERJ (Fundação de Amparo à Pesquisa do Rio de Janeiro), CNPq (Conselho Nacional de Desenvolvimento Científico e Tecnológico). Funding bodies were not involved in study design, data collection, analysis, interpretation, or writing of the manuscript. P.C.B.B was supported by FAPESP (Fundação de Amparo à Pesquisa de São Paulo) Grant 2018/16748-8 and 2016/02978-6, and CNPq for scholarships. F.J.B. acknowledges a Science Without Borders Visiting Senior Scientist Scholarship, Federative Republic of Brazil, that enabled several visits to the laboratory of W.d.S. at the Institute of Biophysics Carlos Chagas Filho, Federal University of Rio de Janeiro, RJ, Brazil.

---

**Supplemental Figure.**
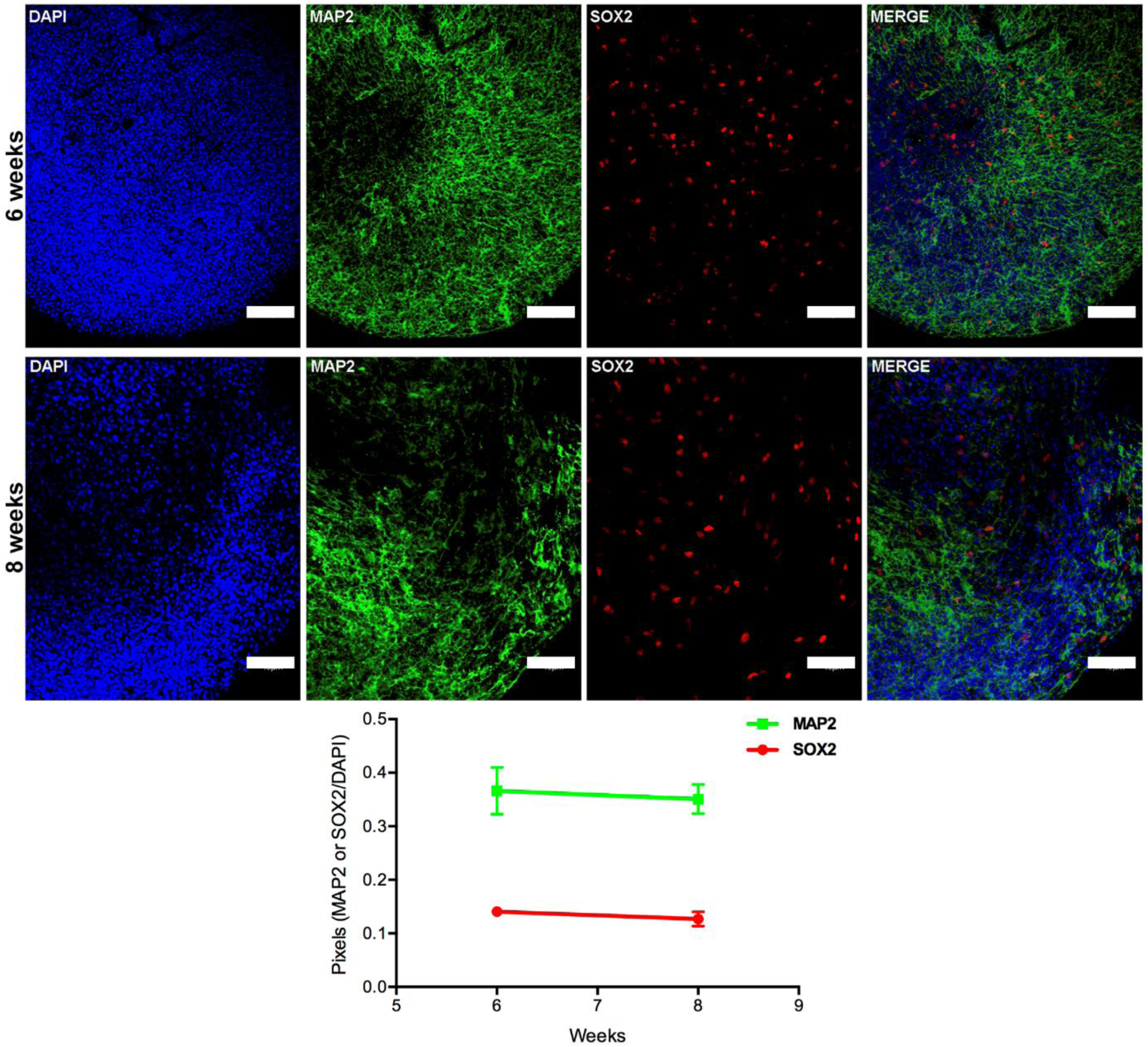
BrainSpheres differentiated from human NPC. Images of BrainSpheres after 6 and 8 weeks of differentiation expressing mature neuron (MAP2, green) and NPC (SOX2, red) markers. Nuclei were stained with DAPI (blue). Scale bars 50 μm with higher magnification. Correlation statistics method revealed a coefficient of variation of 7.25% for SOX2 and 3.04% for MAP2 from week 6 to 8.

